# The Voices of Medical Education Science: Describing the Published Landscape

**DOI:** 10.1101/2022.02.10.479930

**Authors:** Lauren A. Maggio, Joseph A. Costello, Anton Ninkov, Jason R. Frank, Anthony R. Artino

## Abstract

**Introduction:** Medical education has been described as a dynamic and growing field, driven in part by its unique body of scholarship. The voices of authors who publish medical education literature have a powerful impact on the discourses of the community. While there have been numerous studies looking at aspects of this literature, there has been no comprehensive view of recent publications.

**Method:** The authors conducted a bibliometric analysis of all articles published in 24 medical education journals published between 2000-2020 to identify article characteristics, with an emphasis on author gender, geographic location, and institutional affiliation. This study replicates and greatly expands on two previous investigations by examining all articles published in these core medical education journals.

**Results:** The journals published 37,263 articles with the majority of articles published in 2020 (n=3,957, 10.7%) and the least in 2000 (n=711, 1.9%) representing a 456.5% increase. The articles were authored by 139,325 authors of which 62,708 were unique. Males were more prevalent across all authorship positions (n=62,828; 55.7%) than females (n=49,975; 44.3%). Authors listed 154 country affiliations with the United States (n=42,236, 40.4%), United Kingdom (n=12,967, 12.4%), and Canada (n=10,481, 10.0%) most represented. Ninety-three countries (60.4%) were low- or middle-income countries accounting for 9,684 (9.3%) author positions. Few articles were written by multinational teams (n=3,765; 16.2%). Authors listed affiliations with 4,372 unique institutions. Across all author positions, 48,189 authors (46.1%) were affiliated with a top 200 institution, as ranked by the Times Higher Education ranking.

**Discussion:** There is a relative imbalance of author voices in medical education. If the field values a diversity of perspectives, there is considerable opportunity for improvement.

## Introduction

Academic publishing is the primary means of disseminating scientific knowledge in medical education, and the researchers who author those publications drive scientific advancement. However, recent research across scientific disciplines suggests that author diversity, as measured by gender and race,^1,2^ as well as international representation, as measured by country of origin and institutional affiliation,^3,4^ are lacking. This lack of diversity and representation is important because, as many scholars have argued, more diverse teams do better science.^5,6^ For example, scientists with different perspectives often ask different questions, and different questions can lead to novel insights that propel science forward. What is more, modern science is a team sport, as evidenced by the expanding size of research and authorship teams over the past 20 years.^7^ Therefore, today’s research enterprise is largely a group problem-solving activity where diversity of perspectives is a key aspect of creativity, innovation, and overall excellence.^5,6^

In 2021, Maggio and colleagues examined author diversity and representation through two bibliometric studies describing 963 knowledge syntheses appearing in 14 medical education journals between 1999-2019.^4,7^ In these two studies, we attempted to describe the medical education community writ large by providing a bibliometric snapshot of the field’s knowledge syntheses, with a focus on publication characteristics and author gender, country affiliation, and institution. The intent of this work was to provide insights into the landscape of knowledge syntheses, with an emphasis on understanding the authors whose voices are shaping the medical education discourse and evidence. We found that across all authorship positions there is a similar ratio of men and women; however, when considering country and institutional affiliation, there is an overrepresentation of authors from North America and Western Europe working in highly-ranked institutions.^4^ While eye opening, these studies are limited to a single publication type: knowledge syntheses. And thus, we do not know if these findings mirror the broader landscape of medical education publishing, which has implications for understanding which researchers’ voices are most present in our scientific discourse.

Our previous work built on previous studies that have examined slices of medical education literature using bibliometric methods.^8-12^ For example, in 2013 Lee and colleagues investigated the evolution of medical education publications over a 50-year period that were identified as “medical education” based on search terms.^8^ Three years later, Azer produced a ranking of medical education articles from 13 journals based on citations,^9^ and more recently Madden et. al. described the gender of authors and editors in four medical education journals.^12^ These studies each provide a valuable glimpse into the field’s evidence base; however, we are unaware of any research that provides a current, comprehensive overview of these topics in medical education.

The purpose of this study was to replicate and expand on our previous bibliometric work^4,7^ by describing the authorship landscape of the last 20 years of medical education literature, using the largest database of scholarship assembled to date. In the present study, we examined all articles published between 2000 and 2020 in the Medical Education Journals-24 (MEJ-24) to identify article characteristics, with an emphasis on author gender, geographic location, and institutional affiliation.

## Methods

We conducted a bibliometric analysis of all articles published in 24 medical education journals published between 2000-2020, thereby replicating, combining, and expanding the approach used in our previous work.^4,7^

Our sample consisted of articles published between 2000-2020 in the 24 journals featured on the MEJ-24 (See Appendix for journal listing).^13^ The MEJ-24 has been proposed as “a seed set of journals” that constitutes the field of medical education and was developed by our team using a bibliometric co-citation approach.^13^ For these 24 journals, on August, 27, 2021, we retrieved metadata for 22 of these journals from the database Web of Science (WoS). On the same day, for the remaining two journals not indexed in WoS (*Journal of Graduate Medical Education (JGME)* and *The Canadian Medical Education Journal (CMEJ*), we downloaded article metadata from the Crossref REST API. For all 24 journals, we downloaded the following: journal name, author names and affiliations, publication date, open access status, author keywords, and funding details. We selected WoS based on its well-defined metadata and long history as a valuable tool for bibliometric analyses across multiple disciplines.^14^ For the 22 journals indexed in WoS, we also downloaded the number of times an article was cited, its cited references, and the references of any articles that had cited it. These citation data were unavailable for *JGME* and *CMEJ*. All metadata was organized in an Excel spreadsheet.^15^

From each article, we extracted the names of all authors. However, in order to accurately analyze author-level data it is essential to adequately disambiguate author names that represent the same (or different) people.^16^ Thus, we created a thesaurus of author names to reconcile name permutations (e.g., Artino, AR, Jr and Artino, Anthony R, Jr were both reconciled to Artino, Anthony). Complete details of author name disambiguation and the full author thesaurus are freely accessible on Zenodo.^17^ Once the author data was cleaned, to make a prediction of the authors’ gender using their first name, we utilized the tool Genderize.io.^18^ We recognize that our effort to predict gender is an oversimplification of a complex social construct, especially because an individual’s gender is best described by that individual, and our method did not allow us to capture authors who identify as non-binary. However, there currently does not exist a resource that provides gender information for individual authors. Therefore, we relied on the Genderize.io tool, in keeping with work done previously by authors using bibliometric methods.^4,11,19^ Genderize.io bases its predictions on a database of over 100 million names, and for each name provides the “percent confidence” that the gender prediction is correct. We accepted the tool’s gender prediction if and when it was over 70% confident of the gender prediction.

For all articles, we identified country and institutional affiliations. To identify these affiliations, we included any article with at least one author-institution affiliation. As institutional affiliations are often variably reported in paper, we created an additional thesaurus of affiliations, which enabled us to accommodate these variations (e.g., Univ Nebraska and Nebraska Univ reconciled to University of Nebraska) and to associate institutions (e.g., university hospitals) with a parent institution. The thesaurus and full details of its creation are available on Zenodo.^17^ To provide context for institutions, we identified each institution’s ranking on The Times Higher Education World Ranking 2022,^20^ which includes over 1,600 institutions worldwide. Additionally, we identified the World Bank Classification,^21^ for each country which is based on their gross domestic product per capita and assigns countries to four regions (low income, lower-middle income, upper middle income, and high income).

We used Excel^15^ to calculate descriptive statistics.

## Results

Across the MEJ-24, 37,263 articles were published. Of the 37,263 articles, 47 listed no author and 212 were anonymously written. We excluded these articles (0.7%) from our analysis given our emphasis on authorship characteristics and to facilitate consistency across analyses. Of the remaining 37,004 articles, the largest number of articles were published in 2020 (n=3,957, 10.7%) and least number in 2000 (n=711, 1.9%). This difference in articles published represents a 456.5% increase over the time period examined. *Academic Medicine* (n=7760, 21%), *Medical Education* (n=5499, 14.9%), and *Medical Teacher* (n=5029, 13.6%) published the most articles, accounting for 49.4% of all publications over this 20-year period in the 24 journals analyzed.

Articles published in the 22 journals with citation data (n=34,652) were cited 548,358 times. On average, articles were cited 15.8 times (SD=45.54 Range:0-3,310). There were 6,464 (18.7%) articles with no citations, of which 1,333 were published in 2020. “Making sense of Cronbach’s alpha” published by the *International Journal of Medical Education* in 2011 was the most cited article in the dataset (n=3,310).^22^ Table 1 lists the top 10 authors by article citations (see Supplemental Table 1 for a list of the top 20 authors).

**Table 1:** Top 10 cited authors publishing in 22 medical education journals between 2000-2020

| **Author (gender)** | **Total citations** | **# Articles published since 2000** | **# Citations for most cited article** | **Average citation rate\*** |
| --- | --- | --- | --- | --- |
| Cees van der Vleuten (male) | 15733 | 407 | 658 | 38.7 |
| Olle ten Cate (male) | 8697 | 215 | 561 | 40.5 |
| Geoff Norman (male) | 8514 | 209 | 1435 | 40.7 |
| David Cook (male) | 7887 | 114 | 1661 | 69.2 |
| Albert Scherpbier (male) | 7149 | 228 | 337 | 31.4 |
| Kevin Eva (male) | 7106 | 176 | 568 | 40.4 |
| William McGaghie (male) | 6993 | 101 | 1845 | 69.2 |
| Glenn Regehr (male) | 6146 | 126 | 568 | 48.8 |
| Eric Holmboe (male) | 5756 | 155 | 384 | 37.1 |
| Karen Mann (female) | 5326 | 68 | 935 | 78.3 |

\*Average Citation Rate = total citations/total articles published

Nearly half of the articles published in all 24 journals (n=16,817; 55.5%) were publicly accessible. For those that were publicly accessible and that also had citation data (n=34,652), the average citation rate was 15.0, which was just below the overall average citation rate.

### Authors

Of the 37,004 articles analyzed, we identified 139,325 authorship positions. The average author team included 3.8 (range 1-80; Median=3, SD=2.6) authors. If we exclude all single authored papers (n=6,884, 18.6%), the average team size increases to 4.4 members. The largest team included 80 authors who conducted a non-randomized, multicenter trial on anti-stigma training (toward patients with mental illness) for medical students.^23^ In 2000, the average team size was 2.7 (range: 2-14; median: 2) authors, and in 2020, the average team size was 4.3 (range: 2-40; media: 4), which represents a 57.3% increase in average author team size.

By disambiguating the 139,325 author names, we identified 62,078 unique authors representing 14,573 unique first names. For these names, we excluded unknown names (n=1,473) and any names with less than a 70% probability of a gender match (male [n=675], female [n=636]) as predicted by Gender.io. This resulted in 5,418 female and 6,532 male first names used for our analysis.

Applied to our unique author list, Gender.io predicted 62,828 males (55.7%) and 49,975 females (44.3%) total. On multi-author teams, 51.7 % of first authors and 60.4% of last authors were males (See Table 2). In addition, males wrote 67.1% of single-authored articles. Multi-authored articles with female first authors (n=12,739) were cited on average 16.3 times (37.7=stdev) in comparison to 17.7 average (54.4=stdev) for males (n=13,632). Single author citation averages for males and females were 10.4 (34.2=stdev) and 8.6 (40.4=stdev), respectively. Figure 1 provides a comparison of author gender over the 20-year time period for first and last authors on multi author articles.

**Table 2:** Predicted author gender for articles published in 24 medical education journals published between 2000-2020.

|  | Male (%) | Female (%) |
| --- | --- | --- |
| All author positions | 62,828 (55.7) | 49,975 (44.3) |
| Single Author | 3,932 (67.1) | 1,925 (32.9) |
| First Author (Team) | 13,632 (51.7) | 12,739 (48.3) |
| Last Author (Team) | 15,807 (60.4) | 10,376 (39.6) |
| Not first or last author (Team) | 33,389 (42.2) | 26,860 (34.0) |

\*We were unable to predict the gender of 3,964 authors (Genderize.io returned: ‘unknown’ (n=1325), male under .70 probability (n=1406), female under .70 probability (n=1233))

**Figure 1:**
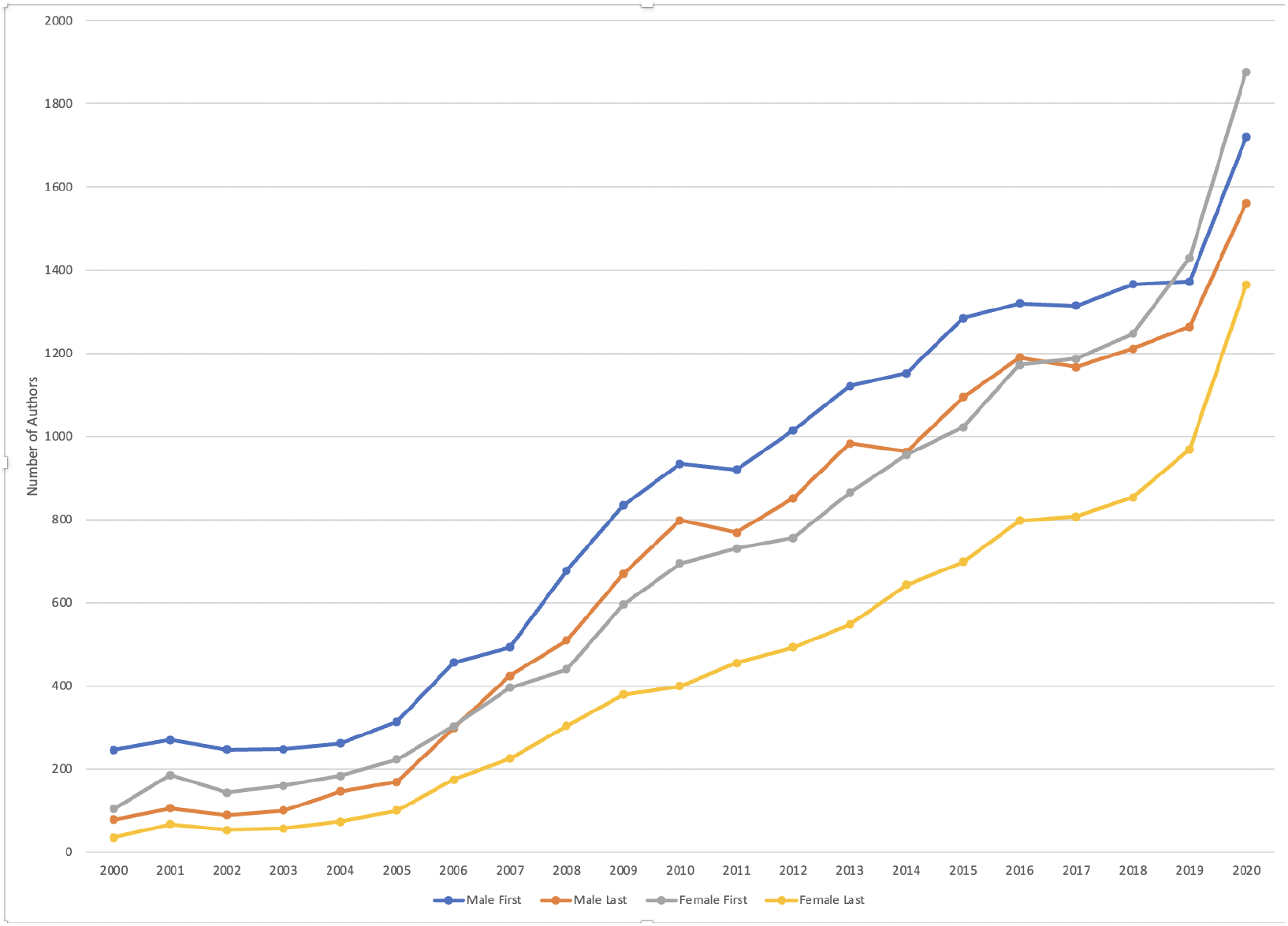
A comparison over time of male and female first and last authors publishing multi-author articles in medical education journals between 2000-2020.

### Geographic Affiliation

Due to limitations in available metadata, we included 28,805 articles with affiliation data from 22 journals in this analysis. Authors listed affiliations for 154 of 189 (81.5%) countries worldwide. The United States (US) (42,236, 40.4%), United Kingdom (UK)(12,967, 12.4%), and Canada (10,505, 10.0%) were most represented country overall, with 69.5% of all articles including at least one author from these three nations. For articles with more than one author (n=23,257), a minority of articles were written by multinational teams (n=3,765; 16.2%). For the 5,617 solo authored articles, authors represented 94 of 189 countries (49.7%), with 381(6.9%) classified as lower and lower-middle income countries (LMIC) (See Table 3).

**Table 3:** Top 10 country affiliations reported in articles published in 22 medical education journals published between 2000-2020.

| Country | Overall appearances of an affiliation | First Author (Team) * | Last Author (Team) * | Single Authors |
| --- | --- | --- | --- | --- |
| United States | 42236 | 8771 | 8377 | 2466 |
| United Kingdom | 12967 | 3263 | 3095 | 1211 |
| Canada | 10505 | 2360 | 2212 | 535 |
| Australia | 5774 | 1373 | 1348 | 271 |
| Netherlands | 5304 | 1009 | 1211 | 154 |
| Germany | 4220 | 803 | 777 | 41 |
| China** | 1786 | 326 | 238 | 29 |
| South Africa | 1358 | 374 | 228 | 97 |
| Saudi Arabia | 1238 | 272 | 156 | 62 |
| Japan | 1013 | 172 | 159 | 12 |

\* Total articles n=23,308 (excludes single author papers (n=5617))
\*\* Lower- and Lower-Middle Income Country

**Table 4:** Top 10 institutional affiliations for authors publishing articles in 22 medical education journals between 2000-2020

| Institution (country) | Overall appearances of an affiliation | Most prolific author (number articles; predicted gender)* | Top cited author (predicted gender)* |
| --- | --- | --- | --- |
| University of Toronto (Canada) | 2476 | Brian Hodges (57;M) | Ryan Brydges (M) |
| Harvard University (USA) | 2380 | Edward Krupat (34;M) | Edward Krupat (M) |
| National Health Service (UK) | 2191 | Paul Baker (22;M) | Ashish Khurana (M) |
| Maastricht University (Netherlands) | 1601 | Cees van der Vleuten (259;M) | Cees van der Vleuten (M) |
| Mayo Clinic (USA) | 1481 | David Cook (102;M) | David Cook (M) |
| University of California, San Francisco (USA) | 1296 | Bridget O'Brien (68;F) | Bridget O'Brien (F) |
| University of Michigan (USA) | 1287 | Larry Gruppen (63;M) | Larry Gruppen (M) |
| Johns Hopkins University (USA) | 1231 | Scott Wright (36;M) | Donlin Long (M) |
| University of Calgary (Canada) | 940 | Kevin McLaughlin (66;M) | Jocelyn Lockyer (F) |
| University of Ottawa (Canada) | 928 | Timothy Wood (47;M) | Stanley Hamstra (M) |

\*Authors matched in thesaurus with matching institution in our thesaurus (n=100,977)

Authors listed affiliations with 93 LMIC accounting for 9,684 (9.3%) author positions. *BMC Medical Education* included the most representation of authors listing LMIC affiliations (n=2,579, 28.1%). For first authors, only 2,552 authors (9.0%) were affiliated with LMIC. (See Table 3 for more details on geographic affiliations)

### Institutional Affiliations

Due to limitations in available metadata, we again included 28,805 articles with institutional affiliation data from 22 journals. We identified 4,371 unique institutions. Of these institutions, 2,112 (48.3%) were degree-granting institutions accounting for 83.2% of affiliations (n=87,057 authors). Alternate institutions included professional associations (e.g., the Association of American Medical Colleges), federal agencies (e.g., United States Department of Veterans Affairs), and hospitals not affiliated with academic institutions (e.g., Alameda Health System, Baillie Henderson Hospital). Of the degree-granting institutions, 957 (21.9%) were ranked by the 2022 THE Ranking: 181 institutions were in the THE top 200 accounting for 48,189 (46.1%) of author affiliations. The top three institutions were University of Toronto (n=2,476), Harvard University (n=2,380), and the National Health Service (UK) (n=2,191). When considering only first and last authors of author teams (n=22,207, n=21,583, respectively), 46.6% of first authors (n=10,573) and 48.0% of last authors (N=10,370) were affiliated with institutions in the THE Top 200.

For multi-author articles (n=23,257), the majority were written by authors from a single institution (n=12,108; 52.1%). The average number of institutions per multi-institutional articles (n=11,150) was 2.9 (range 2-26). The most institutions listed on one article was 26.^24^

## Discussion

This study, which leverages a unique and sizable data set, provides a snapshot of the last two decades of medical education research published in a core set of journals, with a focus on author characteristics. Similar to our previous bibliometric work on knowledge syntheses,^4,7^ we found that the number of medical education articles has grown substantially over the past 20 years. What is more, similar to our previous work, there appears to be a trend towards greater gender parity overall; although male-dominated imbalances in specific authorship positions still persist. Additionally, we observed that the majority of authors list affiliations in Western, English-speaking countries, and these authors tend to be affiliated with highly ranked institutions. Taken together, our findings suggest a relative imbalance of author voices in our medical education science. If diversity of perspectives is important for creativity, innovation, and overall excellence in science,^5,6^ this data indicates considerable room for improvement.

Over the last two decades there was a 456.3% increase in published medical education articles. Although this growth is much less steep than that which we found in published knowledge syntheses (2,620%),^7^ this increase remains impressive and suggests that our field is growing at a relatively fast rate. This increase is likely due to a variety of factors, including the advent of several new journals (e.g., *Focus on Health Professional Education*, which was founded in 2014), a rise in the number of graduate programs in medical and health professions education,^25,26^ and the ever-present pressure to publish felt across many fields.^27^ Looking ahead, it is unclear whether or not this growth trajectory will continue. For example, the journal *BMJ Simulation and Technology Enhanced Learning* ceased publishing articles at the end of 2021.^28^ Additionally, there has been a rise in alternative forms of knowledge dissemination in medical and health professions education (and across science); these alternatives include, for example, podcasts, blogs, and wikis, all of which continue to grow in popularity.^29^

We identified that, overall, there were more males across all authorship positions. However, we also found that the number of female authors is on the rise. Both of these findings generally align with the trends observed in science more broadly,^30^ and specifically in medical education.^4,12^ While these results are encouraging, future work should continue to monitor this trend, especially in relation to the impact of COVID-19. Research conducted to date suggests a downward trend in female authors’ productivity,^31-34^ especially for those with young dependents and those in the early stages of their career.^35^ For example, an analysis of over 40,000 articles published before and during COVID-19 indicated a widening gender gap of 14% percentage points for all authorship positions; that number increased to 24.6% when considering the first author position.^36^

Beyond the basics of author order, our findings suggest that articles solo authored by females, or published with a female as first author of a team, were cited less often than those solo authored by males or published with a male first author, a finding that parallels results from biomedicine.^37^ This finding has implications for the success of women in the field, since citations are often equated with impact and are used to inform high-stakes promotion and funding decisions.^38^ Additionally, fewer female citations in the literature overall indicates that female voices may be less integrated into the field’s evidence-base, which could have important, negative consequences on a field that is increasingly populated by female educators, clinicians, and scientists.^39^

Although author affiliations represented over 80% of the world’s countries, this representation was dominated by authors from English-speaking, Western countries. This finding aligns with previous research in medical education.^4,11^ What is more, only a minority of articles (16.2%) represented multi-national collaborations. This result is somewhat concerning, as articles written by authors from Western countries may fail to recognize non-Western norms and values, which can hinder the translation or adoption of article’s content outside of the local context. While this finding aligns with previous research in medical education, it runs counter to science more broadly, which features a great deal of cross-border collaboration,^40,41^ creating what is considered an “international system of collaboration”.^42^ Adding to the benefits of integrating a variety of international voices, multi-national team research has been positively associated with greater research impact and less research waste.^43^ We propose that medical educators consider approaches to increase international collaborations, which may include future research to understand facilitators and barriers to such collaborations. It might also include the creation of initiatives, such as international funding mechanisms, that encourage cross-border collaboration. It could also include efforts to harmonize research logistics related to data sharing and human subjects protections. For example, in Canada, a recently published consensus statement includes recommendations for researchers to engage in best practices for inter-institutional sharing of medical education data.^44^

Similar to what we found in medical education knowledge syntheses,^4^ we observed in the current study that just under half of authors were affiliated with highly ranked academic institutions, suggesting that these institutions may have a disproportionate level of influence in the field. However, we were encouraged to find contributions from a variety of entities, such as professional associations, federal agencies, and nonprofits. These types of collaborations might help to facilitate knowledge transfer and exchange, and also have the potential to raise the societal value and relevance of research.^45^ As such, these types of collaborations may be worthy of continued growth. That said, in order for authors to reach beyond their national borders and collaborate outside of academia will require support to overcome institutional barriers.^46^ For example, early-career medical education scientists may be less familiar with how to establish and foster collaborations outside of academia. This suggests the need for education on how to negotiate technology transfer and non-disclosure agreements. Currently, such education programs do exist to train academic researchers to partner with individuals outside academia;^47,48^ however, to our knowledge, no such programs exist presently in medical education.

### Limitations

Our findings should be considered in light of several limitations. First, we used the tool Genderize.io to predict author gender based on first names. However, we realize that gender is a complex social construct that is best determined by the individual. Nonetheless, we are unaware of currently available alternatives to establishing author gender in datasets of this scale. Future initiatives might consider encouraging researchers to self-identify their gender in resources such as ORCID. Second, we created our sample based on the MEJ-24, which proposes a seed set of 24 core medical education journals.^13^ While this study expands upon previous efforts by our author team and others to describe a set of core medical education journals,^8,9^ we recognize that our MEJ-24 sample does not include relevant medical education articles published in clinical and other specialty-focused journals. Third, 22 of the included journals publish articles solely in English, with the exception of the *GMS Journal for Medical Education* and the *Canadian Medical Education Journal*, which simultaneously publishes articles in German and French respectively. However, there is some representation from national professional associations such as the *African Journal of Health Professions* and *Focus on Health Professions Education*, which is based in Australia.

Another important limitation is the fact that we were reliant on metadata from CrossRef and WoS, which, by definition, represent a subset of the world’s literature. Additionally, while these databases are often used in bibliometric research, these tools are not infallible and may include data errors that we were unable to detect. In all cases, however, we worked to “clean” our data prior to analysis. The cleaning process required that we create standardized rules. For example, some journals allow authors to list multiple institutional affiliations on the same article. In such instances, which were a minority of cases, we chose to use only the authors first listed institutional affiliation. Notwithstanding this limitation, and in the interest of transparency and the facilitation of future replications, we have provided details for each rule and how it was applied in our dataset.^17^

## Conclusion

Researchers who author medical education articles are the primary drivers of knowledge dissemination and scientific advancement in the field. In this study, we describe the voices of those who have contributed to medical education by examining two decades of the medical education literature, with an emphasis on article and author characteristics. Our findings indicate that medical education scholarship is rapidly growing and evolving, with more female voices being heard, but with a Western country viewpoint that still predominates. Going forward, our community should seek to expand its diversity of voices through collaborations and explicit campaigns that solicit new scholarly perspectives.

## Supporting information

Appendix A: 24 Medical Education Journals

## Ethical Approval

Reported as not applicable

## Data

None reported

## Disclaimer

The views expressed in this article are those of the authors and do not necessarily reflect the official policy or position of the Uniformed Services University of the Health Sciences, Henry M. Jackson Foundation, the Department of Defense, or the U.S. Government.

