## Appendix A: 24 Medical Education Journals for "The Voices of Medical Education Science: Describing the Published Landscape"

### Appendix A: Twenty-four journals identified[1]

*Academic Medicine*

*Advances In Health Sciences Education*

*Advances In Medical Education And Practice*

*African Journal Of Health Professions Education*

*Anatomical Sciences Education*

*BMC Medical Education*

*BMJ Simulation & Technology Enhanced Learning*

*Canadian Medical Education Journal*

*Clinical Teacher*

*Education For Health*

*Focus On Health Professional Education-A Multidisciplinary Journal*

*GMS Journal For Medical Education*

*International Journal Of Medical Education*

*Journal Of Continuing Education In The Health Professions*

*Journal Of Educational Evaluation For Health Professions*

*Journal Of Graduate Medical Education*

*Journal Of Medical Education And Curricular Development*

*Journal Of Surgical Education*

*Medical Education*

*Medical Education Online*

*Medical Teacher*

*Perspectives On Medical Education*

*Simulation In Healthcare-Journal Of The Society For Simulation In Healthcare*

*Teaching And Learning In Medicine*
